# ProbAnnoWeb and ProbAnnoPy: probabilistic annotation and gap-filling of metabolic reconstructions

**DOI:** 10.1101/151258

**Authors:** Brendan King, Terry Farrah, Matthew Richards, Michael Mundy, Evangelos Simeonidis, Nathan D. Price

**Affiliations:** Institute for Systems Biology, 401 Terry Avenue North, Seattle, WA, 98102, USA; Center for Individualized Medicine, Mayo Clinic, 200 First St. SW, Rochester 55905, MN, USA.

## Abstract

**Summary:** Gap-filling is a necessary step to produce quality genome-scale metabolic reconstructions capable of flux-balance simulation. Most available gap-filling tools use an organism-agnostic approach, where reactions are selected from a database to fill gaps without consideration of the target organism. Conversely, our likelihood based gap-filling with probabilistic annotations selects candidate reactions based on a likelihood score derived specifically from the target organism’s genome. Here, we present two new implementations of probabilistic annotation and likelihood based gap-filling: a web service called ProbAnnoWeb, and a standalone python package called ProbAnnoPy.

**Availability and Implementation:** Our tools are available as a web service with no installation needed (ProbAnnoWeb), available at http://probannoweb.systemsbiology.net, and as a local python package implementation (ProbAnnoPy), available for download at http://github.com/PriceLab/probannopy.

**Contact:** http://Evangelos.Simeonidis@systemsbiology.org; http://Nathan.Price@systemsbiology.org

## 1 Introduction

Metabolic modeling approaches provide powerful analytical tools for exploration and detailed consideration of the structure and design of a metabolic network (Schilling et al. 1999). Genome-scale models (GEMs) in particular, which collect all available metabolic knowledge on a particular organism, have been constructed for an expanding array of organisms based on annotated genome sequences (King et al. 2016). GEMs have applications in metabolic engineering, modeling of microbial communities, and simulations that combine transcriptomics, proteomics, and/or metabolomics to deepen understanding of an organism’s phenotype (Milne et al. 2009; Oberhardt et al. 2009).

When a model is not readily available for an organism, or when existing models are not detailed enough to cover the required elements of metabolism for the intended analysis, a new reconstruction needs to be built. Metabolic reconstruction is a data intensive but well defined process (Thiele and Palsson 2010) that requires collecting species-specific information from genome annotations, high-throughput experiments, the literature, and publically available databases, such as KEGG (Kanehisa et al. 2008) or EcoCyc (Karp et al. 2005). Gap-filling methods (Reed et al. 2006) are subsequently applied to improve connectivity to the point where the model can simulate steady state reaction flux and growth.

Most gap-filling tools use an organism-agnostic approach; one that does not consider the relationship between genome and metabolism in selecting candidate reactions from a database. One such example is parsimonious gap-filling, which fills the model using a universal database such as Model-SEED with as few reactions as possible (Devoid et al. 2013). Conversely, our likelihood based gap-filling (Benedict et al. 2014) uses probabilistic annotation to compute organism-specific reaction likelihoods of gene functions based on sequence homology with a trusted annotation database. These likelihoods can subsequently be used to select gap-filling reactions from a biochemical reaction database (Fig. 1). These annotations can additionally provide insight into an organism’s metabolic capabilities and be used in other down-stream modeling tasks.

Here, we provide two new implementations of our annotation likelihood algorithm and its application to likelihood based gap-filling: ProbAnnoWeb, a web service, and ProbAnnoPy, a downloadable python package. Our tools are compatible with openCOBRA packages for con-straint-based reconstruction and analysis (Schellenberger et al. 2011). Mackinac (Mundy et al. 2017), a recent tool bridging COBRApy and ModelSEED, additionally provides functions for generating reaction likelihoods and gap-filling models within ModelSEED, as well as transferring ModelSEED models to and from COBRApy. We propose our tools as an accessible, easy to use, standalone version of probabilistic annotation, with direct integration with COBRApy for likelihood based gap-filling, and as a minimalist alternative for those who wish to compute locally or to have less interaction with online modeling services.

## 2 Methods

### 2.1 Probabilistic Annotation and Reaction Likelihoods

Given an organism’s genome sequence, probabilistic annotation assigns an organism-specific likelihood score (0 ≤ *s* ≤ 1) to each reaction in a template model database of reactions, which comprises the complete pool of candidate reactions for the gap-filling problem. Below we provide a quick overview of this process, which can be explored in greater detail in (Benedict et al. 2014).

First, we run BLASTp on each gene in the query genome against a reference set of high confidence functional annotations (Altschul et al. 1990; Camacho et al. 2009). A log score for each query/target gene pair is computed as follows: 
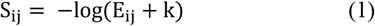
where *E*_*ij*_ is the BLASTp E-value of the pair and *k* is a small constant (10^-200^).The probability that a query gene *i* is in the set of genes *A*_*a*_ with functional annotation *a* is proportional to the score between query *i* and each reference target *j* in *A*_*a*_: 
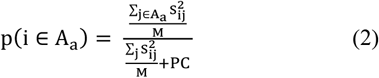
where *M* is the maximum log-score of any BLASTp hits and *PC* is a pseudo-count that dilutes likelihoods for annotations with weak homology to the query.

A reaction’s likelihood is a function of its corresponding annotation likelihoods derived from its Gene-Protein-Reaction relationship specified in a ModelSEED (Overbeek et al. 2005) template model. For iso-enzymes (i.e. “OR” relationships) we take the maximum of enzyme likelihoods, whereas for multi-enzyme complexes (i.e. “AND” relationships) we take the minimum.

### 2.2 Probabilistic Annotation and Gap-filling

Gap-filling can be formulated as a mixed integer linear programming (MILP) problem: the reactions in the model are considered in union with those in a universal template model, a non-zero or minimum increase constraint is placed on the model’s objective function, and the count of new reactions carrying non-zero flux is minimized. In parsimonious gap-filling, each reaction (*x*) not found in the model receives a gap-filling objective coefficient of one (λ_*gapfill,x*_ = 1), and each reaction in the model receives a coefficient of zero.

Likelihood based gap-filling re-weighs the objective coefficients for database reactions according to likelihood as follows: 
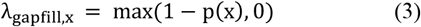

**Fig. 1.**
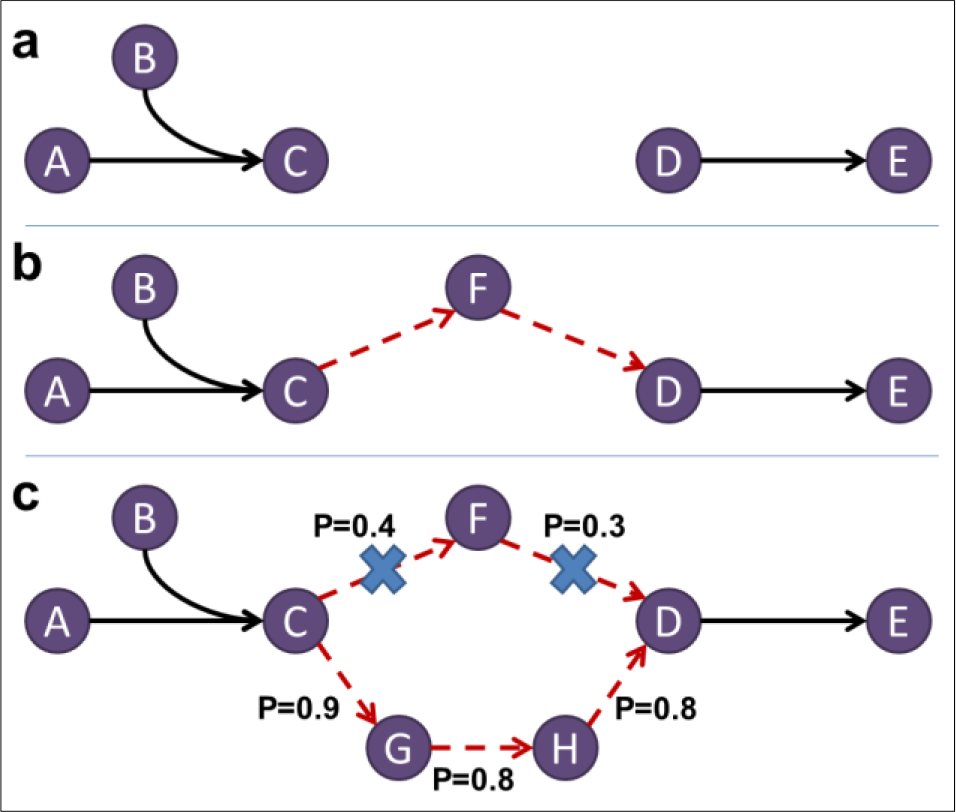
Likelihood based gap-filling. A. Gap in the metabolic network, preventing production of E; b. A parsimonious gap-filling solution using two new reactions; c. A likelihood based gap-filling solution using three new reactions, which are together more likely than the parsimonious solution.

As such, solutions composed of higher likelihood reactions become more favorable, despite potentially requiring more reactions than the optimal parsimonious solution.

## 3 Workflow

We make two tools available: ProbAnnoWeb, a web service, and ProbAnnoPy, an installable python package. The latter is compatible with Python 2.7, models in COBRApy (Ebrahim et al. 2013) format, and depends on usearch, which is freely available for academic use (Edgar 2010). Like COBRApy, ProbAnnoPy depends on an installed MILP solver such as Gurobi. Greater detail on dependencies and the installation process is available at our GitHub repository.

### 3.1 Generating Reaction Likelihoods

In the general use case, the workflow begins with finding a genome sequence for a target organism by downloading its proteome sequence in FASTA format. Next, we run probabilistic annotation, which takes the FASTA sequence and a template model as arguments. The template models serve as general databases for reactions and come from ModelSEED. We supply template models for “Gram Positive”, “Gram Negative”, and “Microbial” organisms. Template choice does not affect likelihoods, only the reactions for which scores are calculated. Probabilistic annotation returns likelihoods in the form of a ‘ReactionProbabilities’ object, which is a wrapper for a dictionary of reaction likelihoods and other information, such as complex and annotation likelihoods.

### 3.2 Likelihood based Gap-filling

We provide functionality for likelihood based gap-filling given a model in COBRApy format, a choice of “universal” model to serve as reaction database, and reaction likelihoods. We support functionality for building a “universal” model from one of the supplied template models, a step that is automated behind the scenes in the web service. Although COBRA isidentifier-agnostic, our implementations of probabilistic annotation use ModelSEED identifiers; currently, only these identifiers are directly supported for probabilistic annotation and likelihood based gap-filling. Like COBRApy’s native parsimonious gap-filling, we output a list of reactions that can be added to the model. ProbAnnoWeb additionally automates this step, instead outputting a gap-filled model that can be downloaded in SBML format.

## 4 Discussion

Probabilistic annotation is a useful tool both for the analysis of metabolic networks and for likelihood based gap-filling, resulting in higher quality reconstructions corresponding to more genomic evidence. Here, we make freely available to the community a straightforward implementation of this algorithm, which can be used to gap-fill metabolic models with ModelSEED reactions. We provide multiple interfaces to our implementation for varied technical needs and levels of programming savvy. Further work will extend applications of probabilistic annotation and support alternative identifier paradigms.

## Acknowledgements

The authors thank Nicholas Chia for important discussions and support.

### Funding

This work was supported by the United States Department of Energy's Advanced Research Projects Agency-Energy [grant number DE-AR0000426 to N.D.P.] and the Mayo Clinic Center for Individualized Medicine [M.M.].

*Conflict of Interest:* none declared.

